# Good host - bad host: molecular and evolutionary basis for survival, its failure, and virulence factors of the zoonotic nematode *Anisakis pegreffii*

**DOI:** 10.1101/2021.03.12.435207

**Authors:** Željka Trumbić, Jerko Hrabar, Nikola Palevich, Vincenzo Carbone, Ivona Mladineo

**Affiliations:** University Department of Marine Studies, University of Split, 21000 Split, Croatia; Laboratory of Aquaculture, Institute of Oceanography & Fisheries, 21000 Split, Croatia; AgResearch Limited, Grasslands Research Centre, Palmerston North, 4410, New Zealand; Laboratory of Functional Helminthology, Institute of Parasitology, Biology Centre of Czech Academy of Science, 37005 Ceske Budejovice, Czech Republic

**Keywords:** accidental host, anisakiasis, *Anisakis* spp., drug targets modelling, paratenic host, transcriptomics

## Abstract

Parasitism is a highly successful life strategy and a driving force in genetic diversity that has evolved many times over. Consequently, parasitic organisms have adopted a rich display of traits associated with survival that guarantees an effective “communication” with the host immunity and a balance with surrounding microbiome. However, gain/loss of hosts along the evolutionary axis represents a complex scenario that as contemporary onlookers, we can observe only after a long time displacement. The zoonotic and monophyletic Anisakidae diverged from its terrestrial sister group Ascarididae 150-250 Ma, although a split from their common ancestral host, a terrestrial amniote, seemingly happened already in Early Carboniferous (360.47 Ma). Faced with the sea-level rise during the Permian-Triassic extinction (215 Ma), anisakids acquired a semiaquatic tetrapod host, and as a result of lateral host-switches in Cenozoic, colonised marine mammals, co-evolving with their “new hosts”. Although contemporary anisakids have lost the ability to propagate in terrestrial hosts, they can survive for a limited time in humans. To scrutinize anisakid versatility to infect evolutionary-distant host, we performed transcriptomic profiling of larvae infecting the accidental host (rat) and compared it to that of larvae infecting an evolutionary-familiar, paratenic host (fish). Identified differences and the modeling of handful of shared transcripts, provides the first insights into evolution of larval nematode virulence, warranting further investigation of shared transcript as potential drug therapy targets. Our findings have also revealed some key intrinsic cues that direct larval fate during infection.

## Introduction

Nematodes from genus *Anisakis* reproduce in marine mammals and propagate embryonated eggs via cetaceans faeces into marine environment, where hatched or in egg-moulted larvae (L1 and L2, or first- and second-stage larvae) use intermediate (copepods and euphausiids) and paratenic (fish, cephalopods) hosts to moult into infective L3 (third-stage larvae), subsequently reaching the gastric chambers of the toothed whales (L4, or fourth-stage larvae, adults) through the trophic chain (see Mattiucci et al, 2018; Mladineo and Hrabar, 2020). In the paratenic hosts, being the most important from the human epidemiology standpoint, larvae perforate gastrointestinal wall and spiralise on visceral surface of abdominal organs in a dormant stage of paratenesis (Beaver, 1969). Therefore, the level of tissues damage in paratenic hosts is usually limited and reversible, although persistent infections and an elevated number of dormant larvae might interfere with its fitness (Levsen et al., 2018). More importantly, infective L3 can also migrate *intra vitam* or *post mortem* into paratenic hosts’ musculature (Cipriani et al. 2016).

Humans are considered only accidental hosts that contract L3 through consumption of inadequately prepared seafood, therefore the associated diseases have mainly been recognised in countries with larger per capita fish consumption (Pozio, 2013). Described for the first time in 1960 in the Netherlands (Van Thiel, 1960), human anisakiasis is a disease caused by infective larvae of genus *Anisakis* spp. (Anisakidae, Nematoda), occurring as gastric, intestinal, ectoptic or gastro-allergic form (Audicana and Kennedy, 2008), or eventually as an asymptomatic form within *Anisakis*-seropositive population (Moneo et al., 2017). Although it shows an ambiguous epidemiological status worldwide (Bao et al., 2019), anisakiasis listed as fifth in the European risk ranking, and the second of 24 foodborne parasitoses with the highest “increasing illness potential” (Bouwknegt et al., 2018). Nonetheless, it is still considered underreported in Europe, showing a large bias between clinical reported cases, i.e. 236 between 2000-2016 (Serrano-Moliner et al., 2018), and the predicted annual number deriving from a quantitative risk assessment model, i.e. between 7,700-8,320 only in Spain (Bao et al., 2017).

The damage afflicted in humans as accidental hosts during larval migration has been extensively reviewed (Audicana and Kennedy, 2008; Hochberg and Hamer, 2010), and rodent models mimicking humans as host-type have been used to characterise *Anisakis* sp. pathogenicity and virulence (Romero et al., 2013; Zuloaga et al., 2013; Lee et al., 2017; Bušelić et al., 2018; Corcuera et al., 2018; Hrabar et al., 2019) (excluding sensitization/ allergy studies). In experimentally *Anisakis-infected* rats, the early response consists of a strong immune reaction with marked induction of specific proinflammatory cytokines and alarmins or damage associated molecular patterns (DAMPS; calprotectins S100A8 and S100A9), regulated through expression of leukocytes-silencing miRNA (*mi*RNA-451 and miRNA-223) (Bušelić et al., 2018; Hrabar et al., 2019). Conversely, *Anisakis* experimentally infecting fish induce no significant regulation of cytokines (IL-1β, IL-4/IL-13, IL-6, IL-8, IL-10, IL-22, TNF□ and TGFβ) and downregulation of IgM and CD8 (cytotoxic T cells), suggesting that larval excretory/ secretory products (ESPs) contribute to the immune silencing in this particular host (Haarder et al., 2013). The only significant early upregulation of serum amyloid A (SAA) in liver, an acute-phase protein, evidences for a systematic proinflammatory reaction that subsides by time. This may be the cause of spontaneous resolving of *Anisakis* infections in fish under the experimental conditions, where only few larvae successfully fulfil their role within their natural paratenic host (Quiazon et al., 2011; Haarder et al., 2013; Marino et al., 2013). However, this greatly contradicts the epidemiological omnipresence of anisakids in fish in nature (Levsen et al., 2018).

We hypothesize that each particular host-type, i.e. paratenic and accidental, represents a distinct ecological niche for the parasite that consequently prompts the infective stage to incur dramatically different strategies, eventually leading to a different degree of propagation success within the host. Such “strategies”, translated through the parasite’s transcriptomics signatures, could be very useful to infer factors of virulence, as well as array of adaptational traits necessary for parasite survival, development and interaction with the host that will allow the parasite to thrive. The features of both the virulence [herein defined as a reduction in host fitness caused by infection (Read, 1994), not to be interchangeably used with pathogenicity, which refers to the ability of an organism to cause disease] and adaptations, depicted as perturbed gene groups and pathways, could be in future used for targeted drug therapy.

Therefore, we aimed to evaluate the early (<32 h) transcriptomic response of *Anisakis pegreffii* migrating and non-migrating L3 during experimental infection of an evolutionary-familiar host (i.e. fish, paratenic host) and evolutionary-distant host (i.e. rat, accidental host and model for human infection). We further assumed that the transcripts significantly changed between migrating and non-migrating L3 and common denominators for both hosts would indicate genes essential for successful larval propagation. Finally, to gain insight into the substrate specificities of the genes deemed important for *Anisakis* virulence, we modelled the tertiary structures using the full-length amino acid sequences of a set of differentially expressed genes that are common in both host-types and inferred the size of their homologous families within available helminth genomes.

## Materials and Methods

### Animal Ethics

All animal care and use protocols were conducted in accordance with ethical standards for animal protection. Experimental infections on rats were performed at the Animal Facility, University of Split (permit number HR-POK-19), approved by the Veterinary and Food Safety Authority, Ministry of Agriculture of the Republic of Croatia (permit number EP 18-2/2016) and Ethics Committee of the University of Split, School of Medicine (permit number 003-08/18-03/0001).

Rats were raised and housed in pairs, in plastic cages with sawdust and corn bedding, food and water ad libitum, temperature 22 ± 1 °C, with a 12 h light/dark cycle (Bušelić et al, 2018). The experimental infections of the European sea bass (*Dicentrarchus labrax*) were performed at the experimental hatchery facilities of the Institute of Oceanography and Fisheries. Fish (n=32) were transported from a nearby farm and transferred to a single concrete flow-through tank (12 m^3^) were they were fed commercial dry formulated diet and kept under natural photoperiod. Water parameters (salinity, temperature and dissolved oxygen) were measured daily by OxyGard H01PST probe. The fish were then distributed into additional four tanks for 15 days acclimatization prior to experimental infections, so that each tank contained seven fish, and the fifth accommodated four fish that served as a negative control.

### Experimental Infections

*Anisakis* spp. larvae were collected from natural sources. Blue whiting *Micromesistius poutassou* were freshly caught and immediately iced by commercial fishermen in the Adriatic Sea and delivered to experimental facilities on the morning of the experiment. Actively moving larvae were carefully removed with forceps from fish viscera and washed in a physiological saline solution. Integrity of their cuticle was checked under a stereomicroscope (Olympus BX 40) and larvae were kept at 4 °C until experimental manipulation, when they were placed in gastric probes. At this point, a sample of pre-infection larvae was preserved in Tri Reagent (Ambion Inc., Invitrogen, Carlsbad, CA, USA) at −80 °C for RNA extractions.

The duration of experimental infections with the two hosts was determined in preliminary experiments. For the detailed description of preliminary findings and the setup of the final experiment on rats, the reader is referred to Bušelić et al. (2018). Briefly, rats were separated in individual cages 24 h prior to the experiment and deprived of food. Thirty-five male Sprague-Dawley rats were used for the final experiment (average weight 207 ± 20.1 g), split into five groups each sacrificed at 6, 10, 18, 24 and 32 h post-infection. Within each group, five rats were intubated with 10 *Anisakis pegreffii* larvae each and two were intubated with 1.5 mL of physiological saline solution serving as external controls. For the intubations, rats were anesthetized using a mixture of 50-100 mg/kg of Ketaminol (Richter Pharma AG, Wels, Austria) and 5-10 mg/kg of Xylapan (Vetoquinol UK Ltd, Buckingham, UK) by intraperitoneal injection. At specific post-infection time-points, animals were administered an overdose of Ketaminol (Richter Pharma AG, Wels, Austria; > 150 mg/kg) and decapitated to confirm death. Following dissection and gross pathological examination, damaged host tissue and recovered *A. pegreffii* larvae were collected and stored in Tri Reagent (Ambion Inc., Invitrogen, Carlsbad, CA, USA) at −80 °C. Prior to conservation, larvae were washed in physiological saline solution to remove all visible traces of host tissue.

Similarly, European sea bass were deprived of food 24 h prior to the experimental intubation with *A. pegreffii* larvae. In total, 32 male sea bass (average weight 92.9 ± 22.2 g, average length 20.8 ± 1.8 cm) were used for this experiment, 28 intubated with L3 larvae and four used as external controls intubated with 2 mL of physiological saline solution. Fish were anesthetized by submersion bath into tricaine methanesulfonate (MS222, Sigma Aldrich, St. Louis, Missouri, United States) solution in seawater (15-30 mg/L) and each intubated with 20 *A. pegreffii* larvae or physiological saline solution. The duration of the experiment was limited to 12 h as we observed a high rate of larvae clearance during a preliminary experiment (data not shown) via passage through digestive tract or regurgitation. Sea bass were euthanized at 3, 6, 9 and 12 h post-infection (7 fish per time-point including controls) by an overdose of MS222. Animals were inspected as previously described for the rat experiment and samples were stored for further analyses.

### Extraction of Nucleic Acids

Total RNA was extracted from total sampled larvae for transcriptomic analyses and DNA was subsequently extracted from reaction leftovers in order to identify *Anisakis* species at molecular level. By doing so we managed to use molecular information from an entire larva for transcriptomics and avoid possible losses due to tissue fragmentation that would be required for separate DNA preparations. Total RNA was extracted using Tri Reagent (Ambion Inc., Invitrogen, Carlsbad, CA, USA) as per manufacturer’s instructions and dissolved in 20-40 μl of RNase/DNase free water (Merck Millipore, Billerica, MA, USA). Quantity, purity and integrity of RNA preparations were assessed by spectrophotometry and 1% agarose gel electrophoresis, respectively. DNA was extracted from Tri Reagent discarded after phase separation for each individual RNA extraction as described in the manual. Molecular identification of larvae was performed using PCR-RFLP method described by D’Amelio et al. (2000). Restriction pattern of rDNA characteristic of *Anisakis simplex* (sensu stricto) × *A. pegreffii* putative hybrid was observed for a single larva and all other were confirmed as *A. pegreffii*.

### Library Preparation for Illumina Sequencing

Samples were sequenced at the Laboratory for Advanced Genomics, Division of Molecular Medicine at the Ruđer Bošković Institute, Zagreb, that also performed cDNA library preparation. Quality and quantity of RNA extractions were checked using a 2100 BioAnalyzer (Agilent Technologies, Santa Clara, CA, USA) and Qubit 3.0 (Thermo Fisher Scientific, Waltham, MA, USA). Based on the sample quality and the experimental outcome, 16 pools of several biological replicates were created for larvae used in rat infections, 8 for migrating larvae, 6 for non-migrating and one pool of naïve pre-infection larvae. We preserved the information about the tissue of larvae recovery within the host (**Supplementary Table 1**). Larvae sampled from sea bass each constituted individual samples with 4 libraries for non-migrating, post-migrating and spiralized larvae and 4 for larvae collected prior to the start of the experiment (**Supplementary Table 1**). cDNA libraries were prepared according to TruSeq Stranded mRNA kit (Illumina, San Diego, CA, USA). The paired-end sequencing (75 bp from each end) was performed on the Illumina NextSeq 500 platform (Illumina, San Diego, CA, USA) over four lanes.

### RNA-Seq Raw Reads Pre-processing

FASTQC v0.11.8 (Andrews, 2018) was used to assess read quality and tailor all subsequent read cleaning steps. Trimmomatic v0.39 (Bolger et al, 2014) was used to remove Illumina adapter sequences, cut first 13 bases from the start of a read due to per base sequence content bias, clip the reads using a sliding window of 4 bases with quality threshold of 20 and remove reads shorter than 30 bases, in that order. The proportion of ribosomal RNA (rRNA) reads in each library was assessed and removed using SortMeRNA v2.1 by comparison against rRNA databases included in the software package (SILVA and RFAM) (Kopylova et al, 2012). Finally, reads were screened for host contamination by mapping against respective host genomes, *Rattus norvegicus* (v6), Ensembl release 98 (https://www.ensembl.org/Rattus_norvegicus/Info/Index) and *Dicentrarchus labrax* (dicLab v1.0c available from http://seabass.mpipz.mpg.de/cgi-bin/hgGateway) using STAR v2.7.1a (Dobin et al, 2013) in 2-pass mapping mode. Reads that failed to map to respective host genomes were used for downstream analyses.

### Transcriptome Assembly and Functional Annotation

Paired-end reads passing quality control were concatenated across all samples into a single set of inputs as recommended by Haas et al (2013) and used to reconstruct a reference transcriptome using Trinity v2.8.6. (Grabherr et al, 2011) with default parameters (*kmer* size 25, minimum contig length 200 nucleotides, strand-specific read orientation set to RF). In order to assess the quality of the initial assembly, we calculated assembly N50 (by TrinityStats.pl script from Trinity), overall read alignment rate using Bowtie2 v2.3.5.1 (Langmead et al, 2018) and examined the number of transcripts that appear to be nearly full-length by comparing assembled transcripts against UniProtKB/Swiss-Prot database (The UniProt Consortium, 2019). For this purpose, blastx algorithm (Altschul et al, 1990) was used as implemented in BLAST+ v2.10.0 (Camacho et al, 2009), with an expectation value (*e*-value) cut-off of 10^-20^. Conserved ortholog content of the reference assembly was assessed using BUSCO v3.0.2 (Waterhouse et al, 2017) in transcriptome mode against metazoa_odb9 (creation date: 2017-02-13). To reduce transcriptome redundancy and concentrate on potential coding regions, CD-HIT-EST v4.8.1. (Li and Godzik, 2006) was first used to cluster sequences on a nucleotide level using similarity threshold of 0.99. Open reading frames (ORFs) were predicted on this set using TransDecoder v5.5.0 (Haas et al, 2013) with a minimum protein length of 80 amino acids. Predicted ORFs were searched against Uniprot/SwissProt database using blastp (e-value cut-off of 10^-5^), while HMMER v3.2.1. (Eddy, 2011) was used to compare peptides against PFAM database v31.0. (El-Gebali et al, 2019) to identify common protein domains. Single best open reading frame (ORF) per transcript was retained using TransDecoder --single_best_only pipeline prioritizing homology hits over ORF length. Redundancy was further reduced in the remaining transcript set by clustering highly similar amino acid sequences with CD-HIT v4.8.1 (Li and Godzik, 2006), using an identity threshold of 1.00. Coding sequences were annotated using blastp (*e*-value cut-off of 10^-5^) against The NCBI Reference Sequence (RefSeq) collection of non-redundant proteins (release96, accessed 15^th^ Dec 2019 at https://www.ncbi.nlm.nih.gov/refSeq), as well as against publicly available proteomes of model and closely related nematodes respectively: *Caenorhabditis elegans* (WBcel235, Ensemble release 98), *A. simplex* (A_simplex_0011_upd) and *Ascaris lumbricoides* (A_lumbricoides_Ecuador_v1_5_4), available from WormBase ParaSite (https://parasite.wormbase.org). Trinotate v3.1.1 (Bryant et al, 2017) annotation suite was used to retrieve GO (Gene Ontology), KEGG (Kyoto Encyclopedia of Genes and Genomes) and eggNOG (evolutionary genealogy of genes: Non-supervised Orthologous Groups) annotations from blast results and collect all functional information into a SQLite database used to filter and sort data as needed.

### Differential expression analysis

Paired-end reads of each sample were mapped to the reference transcriptome as strand specific using Bowtie2 v2.3.5.1 (Langmead et al, 2018) and abundances estimated by RSEM v1.3.1 (Li and Dewey, 2011) with default parameters. Gene level estimated counts were imported into R v3.6.3 (R Core Team, 2020) through tximport package (Soneson et al, 2015). Differential analysis of gene expression was performed using DESeq2 package (Love et al, 2014) for Biocondutor v3.10 (Huber et al, 2015). Prior to statistical testing, low count transcripts (with less than four reads summed across at least four samples) were pre-filtered and removed from the dataset and exploratory data analysis was performed using principal component analyses (PCA) to ensure correct grouping of the samples according to phenotype. Read counts were modelled as following a negative binomial distribution, and a generalized linear model was fitted for each gene with multi-factor design that included host, state (migrating vs non-migrating) and the interaction term as fixed effects. The results were generated for three contrasts in total using the Wald test: migrating vs. non-migrating larvae in rat, migrating vs. non-migrating larvae in sea bass and the difference between the two, i.e. the interaction showing if the regulation was different when considering migrating vs. non-migrating *A. pegreffii* larvae in two distinct hosts. Pre-infection larvae collected directly from fish prior to the experiment were not included in the statistical testing as this was not the primary experimental question. Differentially expressed genes (DEGs) were identified at Benjamini-Hochberg false discovery rate (FDR) < 0.05 without fold change cut-off. Expression profiles for DEGs were explored and visualized using a venn diagram generated by *VennDiagram* (Chen, 2018) and *ggplot2* packages (Wickham, 2016) for R. Five target DEG sequences; ATP-binding cassette transporter *abcb9*, UDP-glucuronosyltransferase *ugt*, aspartic protease *asp6*, leukotriene A4 hydrolase *lkha4* and cytosolic non-specific dipeptidase *cndp2*, common for both hosts and therefore considered as virulence factors, were selected for downstream analyse of gene family evolution and protein modelling.

In order to gain whole-systems understanding of the data, enrichment of GO terms and KEGG pathways via KEGG Orthologue (KO) identifiers within sets of DEGs was calculated using *goseq* package, taking gene length bias into account (Young et al, 2010). Enrichment within specific DEG group was tested against all expressed genes (after removal of low count features) as a background. Terms/pathways were identified as significantly enriched/unenriched at Benjamini-Hochberg false discovery rate (FDR) < 0.05. For this purpose, KAAS (KEGG Automatic Annotation Server) was used to obtain more extensive KO mapping for *Anisakis* putative peptides using SBH method (Moriya et al, 2007). Following enrichment analyses that did not produce conclusive results, the frequency of all DEGs per GO terms and KEGG metabolic and signaling pathways was calculated and inspected next to mean log (base 2)-fold changes (log2FC) per term/pathway.

### Gene Family Evolution

The five target DEG protein sequences common for both hosts were searched against the Wormbase Parasite (Howe et al., 2017) protein BLAST database using the BLASTP version 2.9.0+ (Camacho et al., 2009) search with default settings. Proteomes containing hits with over 70% identity, scores at least 50% of the *A. pegreffii* match and hit sequence lengths at least 75% of the query sequence lengths were downloaded from WormBase. We gathered predicted proteomes of selected Nematoda representing Clades III, V, IV, C, I, Platyhelminthes belonging to Clades Monogenea, Trematoda, Cestoda, Rhabditophora, with *Homo sapiens*, *Mus musculus* and *Danio rerio* as outgroups. To determine homologs of the five target *A. pegreffii* DEGs, we identified single-copy orthologous groups (OGs) in the five proteomes using OrthoFinder v 2.5.2 (Emms et al., 2015), with cluster selection based on at least 75% species present with a single protein in each cluster. The sequences of the proteins in the OGs containing the five target proteins were extracted and searched against the “nr” database using the BLASTP function of diamond version 2.0.6 (Buchfink et al., 2015) with default settings. InterProScan version 5.26-65.0 (Jones et al., 2014) was used to annotate protein domains and GO terms in the extracted proteins. The diamond BLASTP and InterProScan results were merged using OmicsBox version 1.4.11 (BioBam Bioinformatics, 2019). The extracted proteins were subjected to phylogenetic analysis with 1,000 bootstrap replicates for full-coverage data performed using MAFFT (Katoh and Standley, 2014). Also, individual maximum likelihood (ML) inferences for each resulting trimmed gene-cluster were generated to infer the species tree under a multiple-species coalescent model. An evolutionary model was selected automatically for each cluster and visualized in Geneious Prime v.2019.1.3 (Kearse et al., 2012).

### Protein modelling and structural analysis

The Position-Specific Iterative Basic Local Alignment Search Tool (PSI-BLAST) (Altschul et al., 1997) was used to compare the protein sequences associated with the DEGs of interest with the corresponding deposited structures in the Protein Data Bank (PDB). Structural models of the set of DEGs that are common in both host-types were constructed by submitting the obtained amino acid sequences to the I-TASSER server (Yang et al., 2015). The modelled DEGs include: cytosolic non-specific dipeptidase (CNDP2, E.C. 3.4.13.18), leukotriene A-4 hydrolase (LKHA4, E.C. 3.3.2.6), aspartic protease 6 (ASP6, E.C. 3.4.23.3), ATP-binding cassette sub-family B member 9 (ABCB9, E.C. 7.6.2.2) and UDP-glucuronosyltransferase (UGT3, E.C. 2.4.1.17). To assess the quality of the model and residue geometry, TM-scores (a metric for measuring the similarity of two protein structures and fold similarity) (Zhang and Skolnick, 2004) and C-scores (confidence score of the predicted model) all fell within predicted ranges depicting high confidence in our structures. The best model obtained for each DEG was then further validated using Procheck (Laskowski et al., 1996) and ProSA-web (Wiederstein and Sippl, 2007). The active site residues were deduced and visualized using PyMOL Molecular Graphics System v2.0 (Schrodinger, 2010) where the model was superimposed with their parent structures co-crystallized with substrate or inhibitor.

### Data availability

Raw sequence reads were submitted to NCBI Sequence Read Archive (SRA) and are available under BioProject accession PRJNA475982 (runs SRR13870203 - SRR13870233).

## Results

### Description of the experimental outcome

The outcome of experimental infections of Sprague-Dawley rats with *A. pegreffii* L3 larvae have been previously described in Bušelić et al (2018) and Hrabar et al (2019) detailing host tissue damage and response through histopathology, TEM analyses, immunofluorescence screening, miRNA and transcriptomic profiling. Here we focus on the infecting larvae display and briefly recap some of those results as to facilitate the comparison with the experimental outcome in sea bass. In rats, most of the larvae were found passing through the digestive tract (51%), designated as i) non-migrating, as they did not engage in host tissues penetration (**Table 1**); ii) actively penetrating (migrating through) host gastric and intestinal wall or abdominal wall muscles (37.8%). The latter caused mild to severe haemorrhages visible upon gross pathological examination, oedema, inflammatory infiltrate, compression and necrosis. A few larvae were found dwelling in the abdominal cavity presumably after active penetration of host gastrointestinal tract, designated as iii) post-migratory (6.1%), or iv) in the early stages of spiralization, i.e. settling on the serosa of internal organs and peritoneum (5.1%) (**Table 1**).

Larvae showed no synchronized behavior in respect to specific time post-infection and were found at different migratory stages at all sampling points, although the incidence of non-migratory larvae has decreased with increasing time post-infection and the incidence of all migratory phenotypes has accordingly increased (**Table 1**). However, larvae clearance also intensified with the passage of time as median recovery rate was 90% at 6 h post-infection and only 25% at 32 h post-infection (**Table 1**).

In contrast to the outcome observed with the accidental host, experimental infections with sea bass, a paratenic host, demonstrated much larger clearance rate with median recovery of only 12.5% at 3 h post-infection and 0 at 9 h (**Table 1**). Large individual variability was observed for sea bass at each sampling point, with some fish clearing all intubated larvae, hence the time duration of the experiment was reduced and number of intubated larvae per animal increased in respect to rats. The vast majority of larvae were found passing through the digestive tract (67.6%), followed by larvae dwelling in the abdominal cavity or initiating spiralization on the surfaces of liver, stomach, gall bladder or visceral fat (15.5% each) and a single larva (1.4%) was found during active migration penetrating the wall of the small intestine (**Table 1**). In the latter case, no gross pathological signs of the process were observed, except for a minor perforation. This larva was sampled for a purpose different from RNA-seq (Hrabar et al 2019), therefore was not included in this study. All larvae were collected for further analyses, however due to rapid time-course of infection in sea bass and inconspicuous penetration of the larvae, we were not able to collect host tissue samples that would allow us to match the level of investigation we were able to conduct for rat.

The experimental outcomes guided our choice of samples for transcriptome analyses. As most larvae were found actively penetrating rat abdominal muscles, sample pools were created grouping migrating larvae from different post-infection points in respect to those non-migrating, in order to gain robust estimates of *A. pegreffii* larvae gene expression signatures important for the process of initiating host penetration. In sea bass, each library constituted individual larva in different stages of migration (non-migrating, post-migrating, spiralized) as we were not able to construct meaningful pools with the samples obtained.

### Anisakis pegreffii reference transcriptome

In total, 31 *A. pegreffii* samples were sequenced producing on average 28.8 million read pairs per library. On average, 84% of read pairs per sample survived quality filtering steps and a total of 736,718,931 read pairs was used as input for transcriptome reconstruction using Trinity v2.8.6. (Grabherr et al, 2011). Initial de novo transcriptome consisted of 141,685 transcripts with a total of 189,360,713 assembled bases, median contig length of 596 and with N50 value of 2,774 (**Table 2**). Vast majority of input reads was represented by the assembly as their overall alignment rate using Bowtie2 v2.3.5.1 (Langmead et al, 2018) was 99%. A quarter of assembled transcripts (35,704) produced a blastx hit with UniProtKB/Swiss-Prot database. There were 4,016 proteins represented by nearly full-length transcripts, having >80% alignment coverage. The most targeted organism among blast results was *C. elegans* (CAEEL) with 12501 hits. An assessment of conserved ortholog content according to BUSCO recovered 91.8% of near-universal complete orthologs from the 978 in the Metazoa database with a few fragmented and missing (C: 91.8% [S:27.8%, D:64.0%], F:1.0%, M:7.2%, n:978).

After redundancy reduction by clustering of similar sequences and cleaning, final transcriptome was filtered for transcripts with a detectible coding sequence using TransDecoder v5.5.0 (Haas et al, 2013). It was reduced to 36,201 transcripts with predicted ORFs (**Table 2**) that preserved most of the conserved orthologue content of the initial assembly (BUSCO report: C: 90.7% [S:58.5%, D:32.2%], F:1.0%, M:8.3%, n:978). Between 35 and 78% of these sequences were annotated with different public protein databases (**Table 2**), with most hits produced with conspecific *A. simplex* proteome, as expected. Read pairs from each sample were back-mapped to this final transcriptome for differential expression analyses where average overall alignment rate of 88% per sample was achieved. The final reference transcriptome with nucleotide and amino acid sequences as well as functional annotation is available as **Supplementary Table 2**.

### What is specific about A. pegreffii larvae infecting a paratenic and an accidental host?

Overall pattern of gene expression observed in collected *A. pegreffii* L3 larvae during experimental infections of a paratenic, a sea bass, and an accidental host, a rat, is outlined using principal component analyses (PCA) in **Figure 1**. Consistent with our observations during the experiment, there was high variability between samples mirroring unsynchronized behavior of larvae during the experiment that demonstrated no predilection for a site of infection or temporal dynamics of host invasion. The largest source of variance in the data was the host, as samples primarily grouped whether originating from a homeothermic or poikilothermic host, i.e. rat or sea bass/blue whiting, respectively. Of the two, the strength of the response of larvae that managed to infiltrate host abdominal cavity, was much greater in rat than in sea bass (**Figure 1**). A single sample of migrating larvae found penetrating rat intestine presented an outlier going against general trend of variance observed for experimental groups and was removed prior to differential expression analyses.

**Figure 1.**
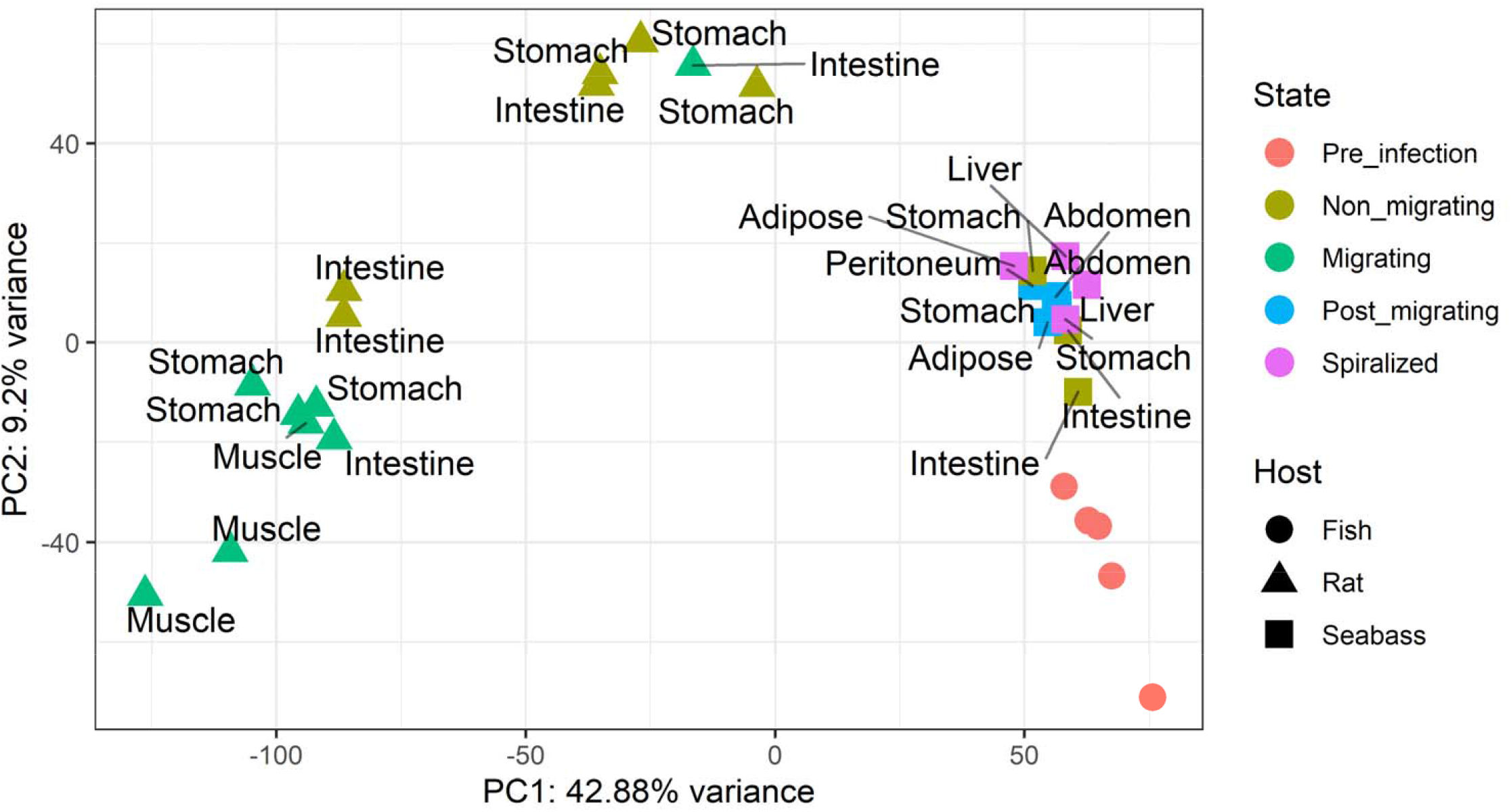
Sample grouping according to gene expression variation in collected *Anisakis pegreffii* L3 larvae during experimental infections of a paratenic and an accidental host, *Dicentrarchus labrax* and *Rattus norvegicus*, respectively. Raw counts were normalized by variance stabilizing transformation and profiles are outlined using principal component analyses (PCA).

Primary experimental question was to unearth general signatures of gene expression paramount for the process of host infection, i.e. those that would be in common to larvae infecting a rat and a sea bass. In turn, delineation of genetic activity that is different between these two hosts might elucidate evolutionary adaptations of *A. pegreffii* important for its survival and propagation through the trophic chain or what is missing when it encounters an unexpected host such as a rat, or human. The statistical design for differential gene expression analyses was constructed with these questions in mind. All samples derived from larvae that passed host barriers were considered migrating and compared against all those that failed to do so within the same host. Pre-infection larvae collected prior to the start of the experiment were not included in the analyses and are shown for exploratory purposes only.

After removal of low count genes, 1,937 were found differentially expressed at FDR < 0.05 in migrating vs. non-migrating larvae in rat (1,096 up and 841 down), 484 in sea bass (328 up and 156 down) and 509 showing evidence of interaction between the main effect and the host, i.e. they were differently regulated in the two hosts. Ten DEGs were at the intersection between all three groups (**Figure 2**) suggesting their importance as contrasting factors in *A. pegreffii* infection of an accidental and a paratenic host. Largest difference in log2FC between migrating larvae in rat and in sea bass was observed for a putative cuticle collagen (CO155) and glucose-6-phosphate exchanger (G6PT1) that were both upregulated in rat and downregulated in sea bass. The reverse was true for NADH-dependent flavin oxidoreductase (NADA) (**Table 3**). Another 65 transcripts were found differentially expressed in migrating larvae in rat and in sea bass showing consistent regulation in both hosts and these are deemed paramount for the process of host invasion in *A. pegreffii*. Of these, 35 that demonstrated at least two-fold change in gene expression in at least one of the hosts are depicted in **Table 3**. There is a group of putative collagen transcripts upregulated in migrating larvae in both hosts, however more strongly so in larvae infacting a rat than a sea bass. Several catalysts and transporters also feature the list showing moderate and congruent upregulation in both hosts, such as cytosolic non-specific dipeptidase (CNDP2), leukotriene A-4 hydrolase (LKHA4), aspartic protease 6 (ASP6), ATP-binding cassette sub-family B member 9 (ABCB9), UDP-glucuronosyltransferase (UGT3), some of which were chosen as potential drug targets. Unfortunately, we were not able to annotate most strongly upregulated DEGs in this list.

**Figure 2.**
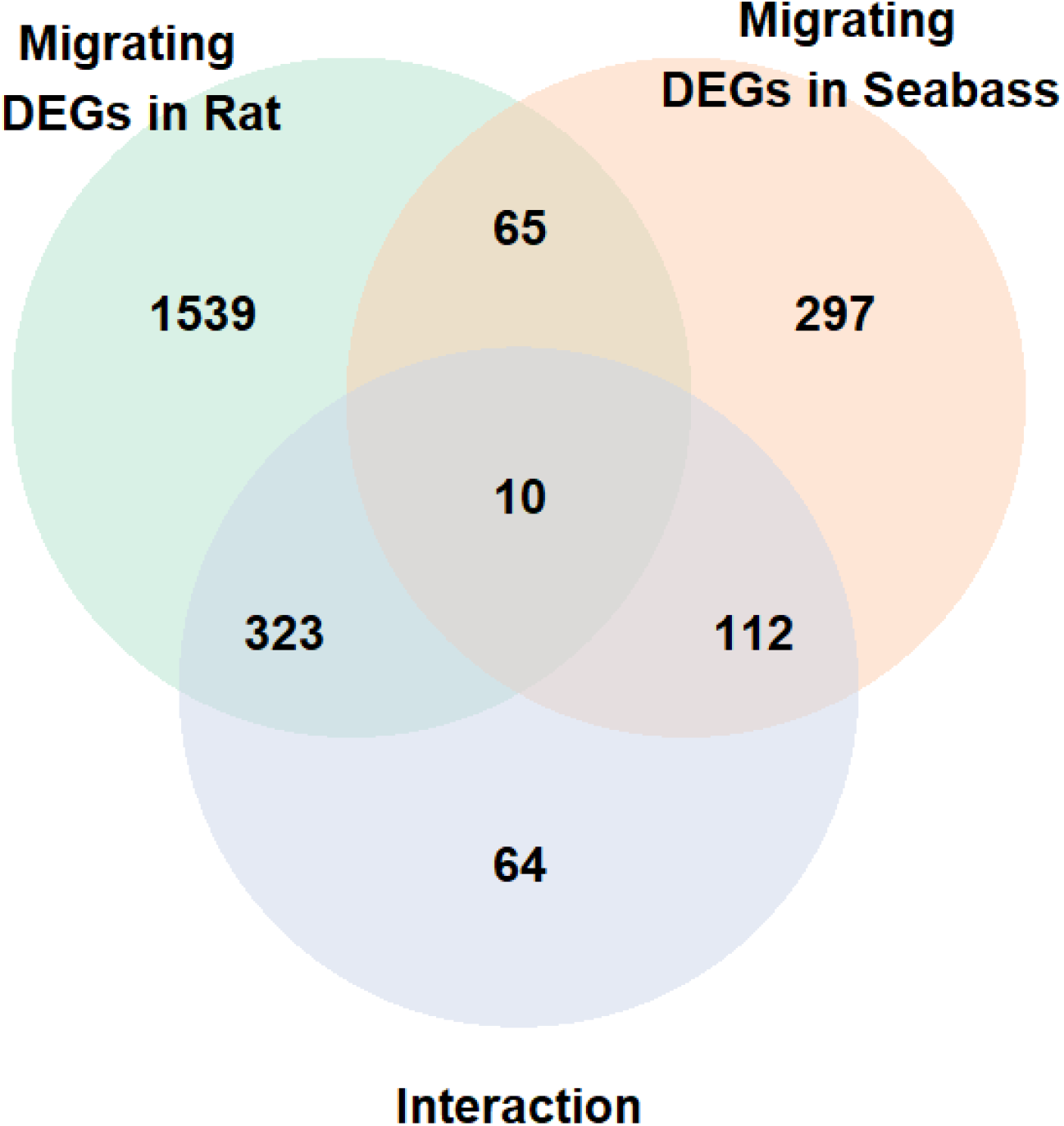
Venn diagram showing overlap between three groups of differentially expressed genes (DEGs) identified in *Anisakis pegreffii* L3 larvae during experimental infections: Migrating vs. non-migrating larvae in rat *Rattus norvegicus*, migrating vs. non-migrating larvae in seabass *Dicentrarchus labrax* and the interaction: larvae showing different regulation in two hosts. All larvae that successfully penetrated host mucosal barriers were considered migrating and compared against all those that remained inside gastrointestinal system of the same host.

Enrichment analyses of GO terms and KEGG pathways resulted in only few significant terms/pathways (FDR < 0.05) for rat DEGs and almost none for other groups (**Supplementary Table 5** and **6**). Over-representation of GO terms: structural constituent of ribosome, structural constituent of cuticle, translation, ATP synthesis coupled proton transport, was found in rat DEGs, as well as enrichment of functions associated with KEGG pathways Ribosome and Oxidative phosphorylation. Since enrichment analyses did not provide conclusive results, to observe DEGs from other groups at a more systemic level we investigated DEG frequency distribution for each GO term and KEGG signaling and metabolic pathway alongside average Log2FoldChange (**Figure 3**). Consistent with the results of enrichment analyses, most upregulated DEGs in rat were counted in KEGG pathways Ribosome, Biosynthesis of secondary metabolites, Oxidative phosphorylation, Microbial metabolism in diverse environments and Carbon metabolism, that also feature as most numerous for sea bass, except for the Ribosome. Two other most represented in sea bass are Lysosome and Autophagy - animal and, although not at the top of the list, there are several DEGs in sea bass associated with drug and xenobiotic metabolism. Most down-regulated DEGs were counted in Endocytosis and Ubiquitin mediated proteolysis in rat, and MAPK signaling pathway and Aminoacyl-tRNA biosynthesis in sea bass. Functional characterization of DEGs based on their frequency in GO categories supports KEGG findings (**Supplementary Figure 1**). Two interesting biological processes grouping several DEGs from rat and sea bass in GO frequency distribution are collagen and cuticulin-based cuticle development and molting cycle showing relatively high average log2FoldChange (>2).

**Figure 3.**
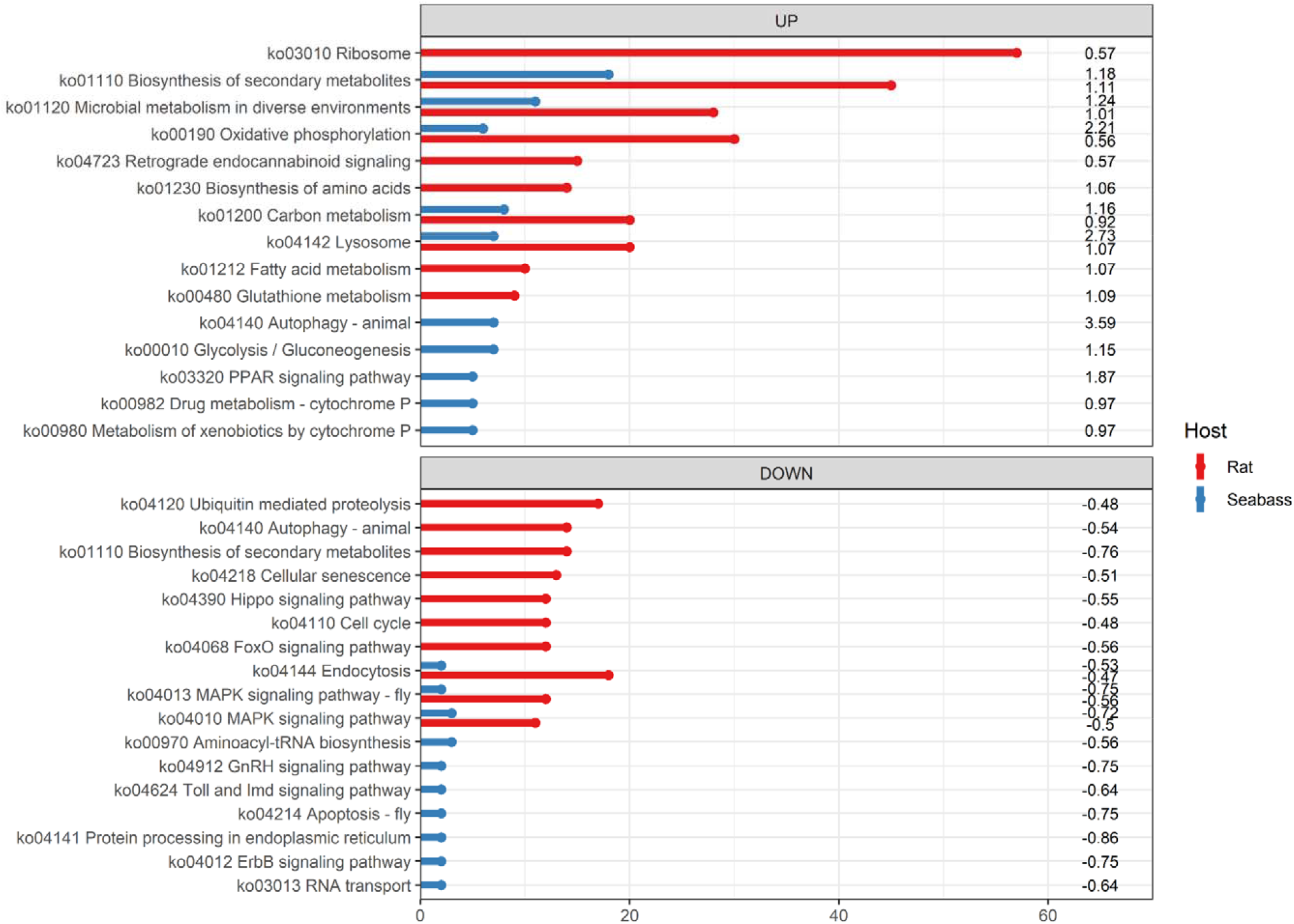
Frequency distribution of differentially expressed genes (DEGs) in top 10 KEGG metabolic and signaling pathways identified in *Anisakis pegreffii* L3 larvae during experimental infections of a paratenic and an accidental host, *Dicentrarchus labrax* and *Rattus norvegicus*, respectively.

Gene-level raw counts used for expression analyses are available in **Supplementary Table 3** and overall results of differential expression analyses for *A. pegreffii* larvae from both hosts are presented in **Supplementary Table 4** were they can be easily filtered to obtain specific DEG groups according to the venn diagram in **Figure 2**.

### Orthology and evolution of significant DEGs

We studied the evolutionary significance of ABCB9, UGT3, ASP6, LKHA4 and CNDP2 proteins in eukaryotes. In particular, we searched the complete genomes or transcriptomes and acquired genome-wide coding sequences from 28 species representing clades I-V of nematoda, platyhelminths and free-living flatworms (monogenea, digenea, cestoda), as well as human, mice, zebrafish as outgroups. For this analysis, we used orthologous groups (OGs) of genes identified based on the five selected and significantly differentially expressed genes of interest for *A. pegreffii* and we modeled gene gain and loss for the five orthologues. Comparative analysis within ABCB9 (OG0000006), UGT3 (OG0001081), ASP6 (OG0000081), LKHA4 (OG0001209) and CNDP2 (OG0000954) revealed numerous highly conserved enzymes present across all nematode clades, but with markedly different orthology profiles (Figure 4). Our phylogenetic analyses indicate that ABCB9s are a broad and ancient eukaryotic gene family, with the only loss reported for *Macrostomum lignano* (Rhabditophora). The UGT3s appear to have been lost independently within eukaryotes, especially in several nematoda (Clades IV and C), Trematoda and Cestoda lineages. In contrast, ASP6s, LKHA4s and CNDP2s appear to be broadly conserved across all species compared in our study, with the exceptions of *Fasciola hepatica* (ASP6), both Clade I nematodes (LKHA4), and *Brugia malayi* (CNDP2). Our preliminary orthology analysis of the ASP6 and ABCB9 transcripts of *A. pegreffii* produced a substantial number of overall homologous sequences (*n*=696 for ABCB9, *n*=296 for ASP6) due to the large and diverse nature of single and multi-domain architectures of eukaryotic ABC transporter/ATP-binding proteins and aspartyl proteases (Supplementary Table 7). To address this, we subdivided clusters of homologous sequences into orthologous groups that matched our *A. pegreffii*transcript annotations and the Enzyme Commission number (E.C.) profiles (Supplementary Tables 2 and 7). In particular, Clade V group nematodes, exhibited massive expansions of ASP6s and UGTs, with tens of homologs that broadly cluster in the phylogeny into two different groups.

**Figure 4.**
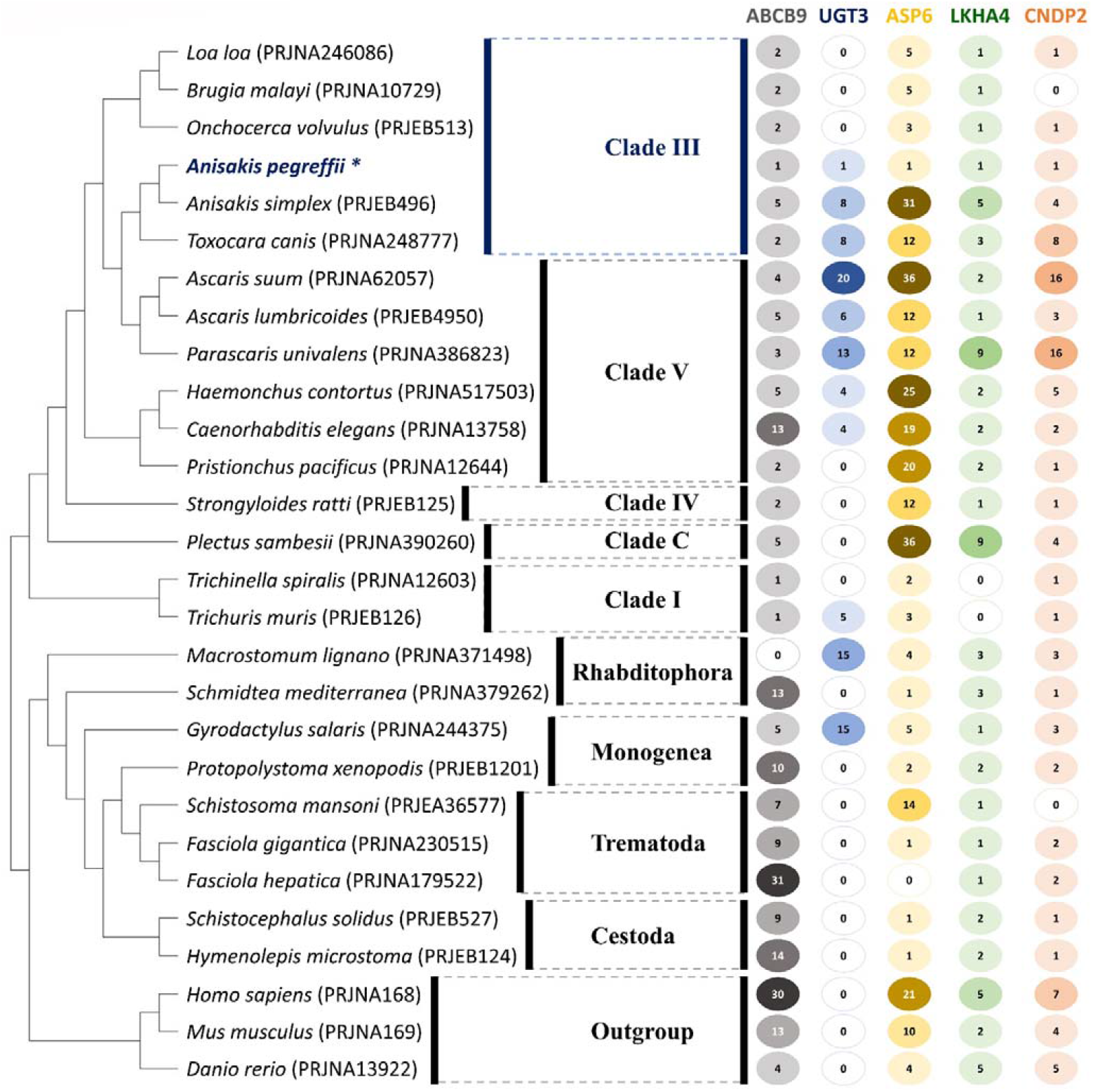
Phylogenetic species-tree reconstruction of homologous gene families corresponding to the five DEGs of interest across 28 species. The consensus tree is based on losses and gains of orthologous groups corresponding to the protein sequence alignments of the five target DEGs of *Anisakis pegreffii*: ABCB9, UGT3, ASP6, LKHA4 and CNDP2 (labelled with *). Only protein sequences that comply within the boundaries of the stringent sequence similarity cut-offs, E.C., GO and InterPro terms and/or descriptions are depicted. The numbers in colored circles represent the total number of gene families corresponding to a particular gene based on orthology. A missing value indicates the absence of an orthologous group corresponding to any of the five target DEGs of *A. pegreffii*. Bioproject GenBank accession numbers are provided (in parentheses) for all reference sequences.

### 3D structure of selected virulence factors and their potential drug-targeting

The five DEGs of interest were modeled using the I-TASSER server to produce tertiary structures of the enzymes (Figure 5). The best of these structures included the ATP-binding cassette transporter ABCB9 (E.C. 7.6.2.2) and the UDP-glucuronosyltransferase UGT3 (E.C. 2.4.1.17; Fig. 5A and C) with C-scores of 1.42 and −1.78 and TM values of 0.54 ± 0.15 and 0.50 ± 0.15 respectively. To further identify the active sites of the enzymes and drug binding potential, these models were superimposed with published crystal structures of structurally homologous enzymes co-crystallised with identified inhibitors. This included 4AYT (Shintre et al., 2013), an ATP-binding cassette (ABC) transporter found in the innermembrane of mitochondria and 6IPB (Zhang et al., 2020), a UDP-glucuronosyltransferase from *Homo sapiens*. The % identity of these sequences with the corresponding *A. pegreffii* sequences were 35.47% and 23.03% respectively. The ABCB9 inhibitor binding site includes residues Tyr-504, Arg-507, Thr-506, Asp-291, Gln-578, Glu-659, Ser-537, Gly-535, Ser-534, Gly-533, Ser-532, Lys-536, Ser-538, Ile-512. It is shown in Fig. 5B bound to the ATP nucleotide analogue phosphomethylphosphonic acid adenylate ester (ACP) that was observed in 4AYT. Overall, this represents an 85.7% sequence identity between active site domains. The active site residues of UGT3 fall within 4 Å the inhibitory tartrate (TTA) molecule observed in 6IPB and includes Ala-309, Phe-310, Gly-311, Asn-312, His-313, Gln-360, Gly-374, His-375, Ala-376, Gly-377, Leu-378, Lys-379, Ser-380, Met-394, Gln-400 (Fig. 5D). Active site identity remains low, (approx. 28.5%), however TTA in both structures makes identical hydrogen bond interaction between the carboxylates moieties and main chain amides of each protein. In fact, few of the observed differences we suspect would preclude TTA binding in our model. For the aspartic protease ASP6, three possible models were produced, however the selected closest structural homologue model had a poor resolution with a C-score of 0.61, a TM value of 0.80 ± 0.09, a RMSD of 3.0 ± 2.2 Å, and normalized z-scores less than 3.48 (Fig. 5E). The protein structure for the ASP6 does not contain the second domain present in the glycoprotein of 6ROW (Scarff et al., 2020) from *Haemonchus contortus* (45.83% identity). The catalytic residues were in part identified via structural comparison with human Progastricsin (1HTR; Moore et al., 1995) and ASP6 and include Gly-53, Thr-54, Ser-55, Phe-56, Asp-71 (Fig. 5F). The selected model for LKHA4 (E.C. 3.3.2.6) had a C-score of 1.42, a TM value of 0.91 ± 0.06 (Fig. 5G) and was modelled after 4GAA (Stsiapanava et al., 2014), a leukotriene A4 hydrolase from *Xenopus laevis*, a bifunctional zinc metalloenzyme co-crystallized with the inhibitor bestatin (BES). Overall, the enzyme shares a 39.09% identity with the *A. pegreffii*, however the active site residues of LKHA4 that fall within 4 Å of BES in our model and the Zn^2+^ binding region shares a 70.5% identity with 4GAA (Fig. 5H). They include Trp313, Met316, Glu320, Tyr388, His301, His297, Leu294, Glu298, Arg560, Leu269, Gly270, Gly271, Met272, Glu273, Phe135, Pro138 and Gln137. For CNDP2 (E.C. 3.4.13.18), a model was produced with a C-score of 1.87, a TM value of 0.98 ± 0.05 (Fig. 5I). The protein structure for the CNDP2 corresponds to 2ZOF (Unno et al., 2008), a carnosinase complexed with Mn^2+^ and a non-hydolyzable substrate analogue bestatin (BES) from *Mus musculus*. The enzyme shares a 54.62% identity with the *A. pegreffii* sequence overall, while the identity of the active site surrounding Bestatin shares a 73.6% identity. The residues of CNDP2 that fall within 4 Å of the inhibitor include Asp-134, His-101, Glu-168, Glu-169, Gly171, His-446, Thr-199, Asp-197, Thr-198, Gln-208, Glu-415, Ile-211, Gly-417, Ser-418, Ile419, Pro420, His-381, Met-214, Arg-344 and His381 (Fig. 5J).

**Figure 5.**
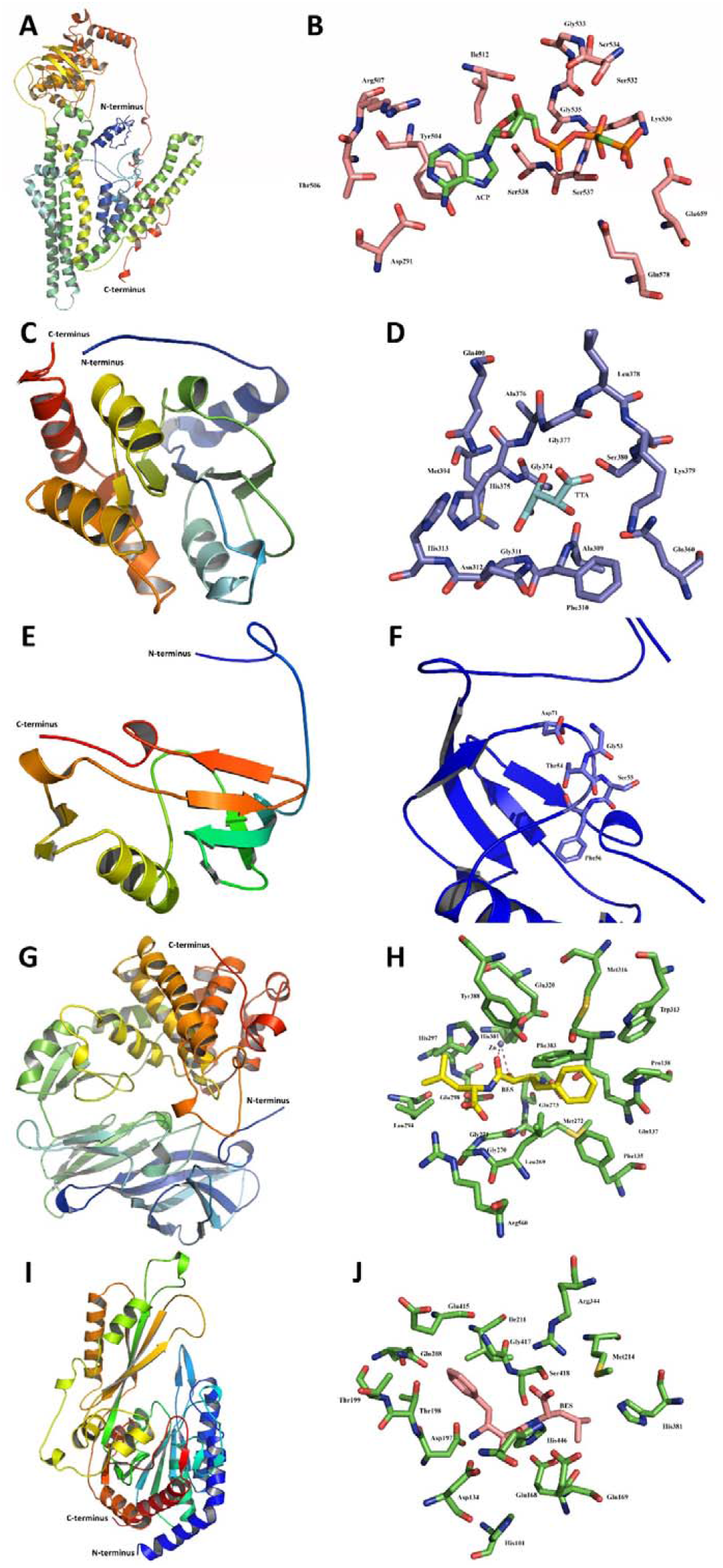
Predicted tertiary structures of the selected drug therapy targets. The analysed DEGs include: A-B, ATP-binding cassette sub-family B member 9 (ABCB9); C-D, UDP-glucuronosyltransferase (UGT3); E-F, aspartic protease 6 (ASP6); G-H, leukotriene A-4 hydrolase (LKHA4); I-J, cytosolic non-specific dipeptidase (CNDP2). Predicted tertiary structures of the five DEG monomers are shown to the right. Locations of the C- and N-terminus in the predicted tertiary structures of each DEG are labelled. The predicted tertiary structure of the ABCB9 (A) showing the location of the active site in salmon (B) and phosphomethylphosphonic acid adenylate ester (ACP) used as an inhibitor (green). The UGT3 predicted tertiary structure (C) with the location of the active site shown in purple (D) and inhibitory tartrate (TTA) in light blue. The ASP6 predicted tertiary structure (E) showing the location of the active site (F) in light blue. The leukotriene A-4 hydrolase (LKHA4) predicted tertiary structure (G) showing the location of the active site shown in green (H) and inhibitory bestatin (BES) in yellow as well as the Zn^2+^ binding region. The cytosolic non-specific dipeptidase (CNDP2) predicted tertiary structure (I) showing the location of the active site shown in green (J) and inhibitory bestatin (BES) shown in salmon.

**Figure 6.**
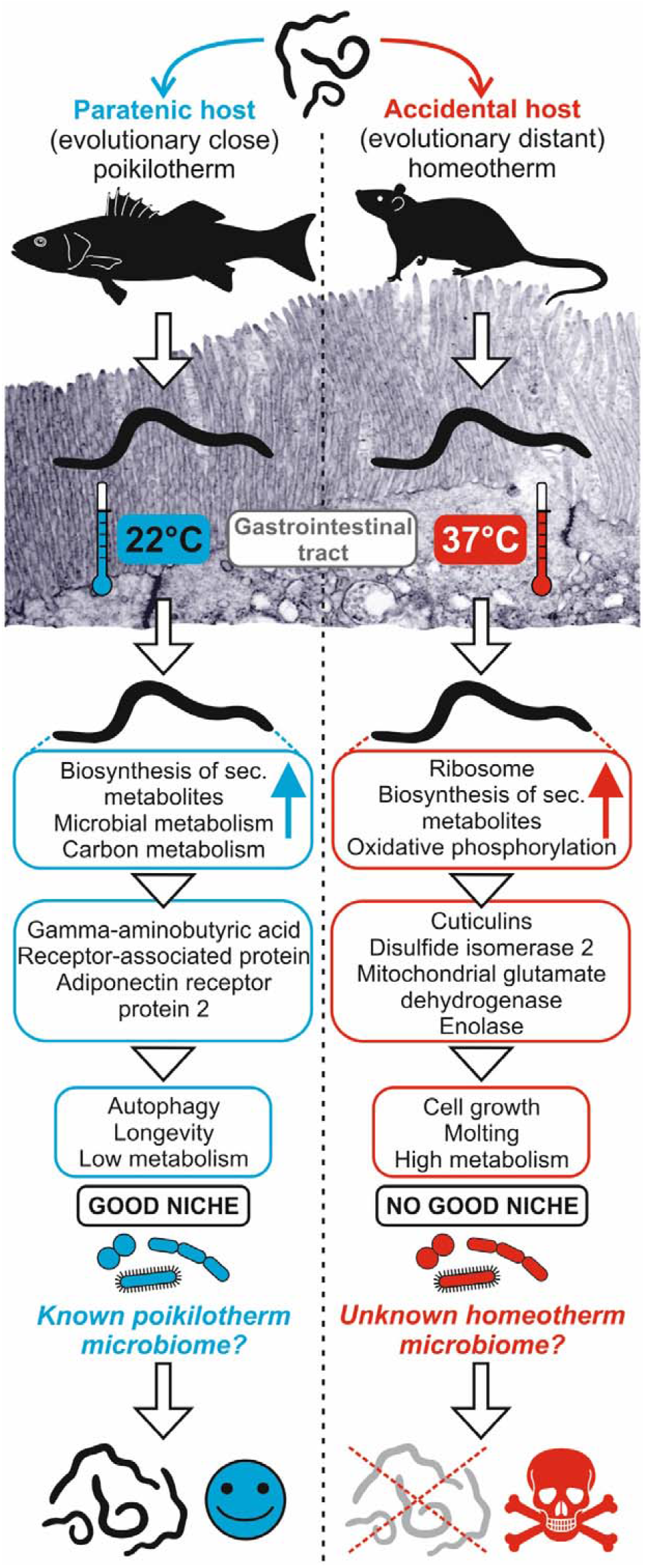
Schematic representation of the fate of *Anisakis pegreffii* L3 larvae after infecting evolutionary-familiar, parathenic host seabass (*Dicentrarchus labrax*) and evolutionary-distant, accidental host rat (*Rattus norvegicus*).

## Discussion

Parasites have a myriad of survival strategies to warrant successful infection and propagation/reproduction within the hosts, being expressed through a highly specific genetic adaptations (Qu et al., 2019). Their genomes show great variability in genome size and organization due to the expansions of non-coding elements, such as long terminal repeat transposons and parasite specific gene families (Coghlan et al, 2019). Despite this, the free-living nematode *Caenorhabditis elegans* still remains the most studied nematode today, building the basis of our understanding of nematode physiology. Functionally, specific gene families in parasites reflect their biology, frequently encoding for proteases/peptidases, protease inhibitors, SCP/TAPS proteins (sperm-coating protein/Tpx-1/Ag5/PR-1/Sc7, from the cysteine-rich secretory protein superfamily), acetylcholinesterases, sensory receptors (G coupled receptors, GPCRs), cuticle maintenance proteins, fatty acid and retinol binding (FAR) proteins, glycosly transferases, chondroitin hydrolase, ABC transporters and kinases, as well as other taxa-restricted gene families involved in niche colonisation and immunity-evasion (Viney, 2018; Coghlan et al, 2019). However, the knowledge on pathways engaged in the active migration of infective larval stages through the host in respect to larvae failing to infect, and the array of responses when larvae face evolutionary-distant host, is mostly fragmented or unsolved for non-model parasites, although it has a practical importance in combating parasitosis.

### How larvae react to good (paratenic) vs bad (accidental) host?

The striking difference between two transcriptome profiles of infecting *Anisakis pegreffii* larvae was largely attributed to host specificity, i.e. larvae infecting the rat (an accidental and evolutionary-distant host) and the fish (a paratenic and evolutionary-familiar host). Less different were profiles in respect to tissues of L3 migration (e.g. stomach, intestine, muscle), or to the designated larval stage (i.e. non-migrating, migrating, post-migrating, spiralised). Expectedly, L3 collected as dormant and spiralised stages from the fish visceral cavity showed profiles divergent from those infecting rat and, to a lesser extent fish, confirming that *Anisakis* spp. persist in paratenic host without essential development and growth (paratenesis) (Beaver 1969). Such evolutionary strategy allows multiple number of encounters between paratenic host and parasite, and consequently, the accumulation and extended lifespan of the latter (Anderson 1982).

Specific life history traits of *Anisakis* are hard to parallel in respect to other parasitic nematodes, creating a gap in knowledge transfer from well-studied and defined laboratory models. Family Anisakidae diverged from its terrestrial sister group Ascarididae approximately 150-250 Ma, but the split from their common ancestral host, a terrestrial amniote, happened already in Early Carboniferous (360.47 Ma). Anisakids acquired a semiaquatic tetrapod host, and as a result of lateral host-switches in Cenozoic, colonised marine mammals and co-evolved with their “new hosts”. Therefore, the most closest referent system is believed to be the intestinal *Ascaris* genus; *Ascaris lumbricoides* that infects humans and causes ascariasis and *A. suum* in pigs (Wang, 2014) [although these are considered a single species based on morphological and genetic similarity, corroborated by the account of cross-infections between humans and pigs (Leles et al, 2012)]. *A. suum* exerts one of the most complex life cycles in its single host; ingested *Ascaris* eggs hatch and the infective L3 larvae undergo an extensive hepatotracheal migration, subsequently returning into the small intestine to reach adult stage. Such behavior somewhat resembles *Anisakis* L3 larvae infecting humans (accidental host), but greatly contrasts the infection pathway in marine mammal, their final host, where no migration occurs. While in *A. summ* each migratory stage exhibits strictly controlled spatiotemporal gene expression patterns, the most abundant transcripts per each stage are shared among stages and can be categorized in three common molecular function GO terms - binding (GO: 0005488; small molecule binding, protein binding, nucleic acid binding and ion binding), structural molecule activity (GO: 0005198; ribosome structural constituent, cuticle synthesis) and catalytic activity (GO: 0003824; hydrolase, oxidoreductase). This coincides well with GOs observed in *A. pegreffii* infecting evolutionary-distant rat and contrasts the observation in the evolutionary-familiar fish described herein. This transcriptomic synergy in *A. summ* and *A. pegreffii* larvae is further confirmed by two top represented KEGG pathways in both parasites; i.e. the Ribosome and Oxidative phosphorylation, the former utterly absent in *A. pegreffii* infecting fish.

A high transcription of ribosome-related genes during infection in homeotherm host reflects the surge in demand for proteins. Whether those involved in building up of larval cellular elements and cell division during growth towards L4 stage, or those necessary for enzymatic reactions related to increased energetic demands and production of essential excretory/ secretory products, can be scrutinised from downstream pathways. The ribosome production during the physiological cell cycle in higher eukaryotes starts at the end of mitosis, increases during G1, is maximal in G2 and stops during prophase (Leung et al., 2004), therefore upregulated Ribosome confirms ongoing of cell proliferation and growth in larval *A. pegreffii* infecting rat. The fact that this is observed only in the accidental hosts where larvae attempt to mature, but failing to reach an adequate attachment niche keep migrating, suggests a higher energetic and metabolic burden imposed on the larvae, eventually resulting in spent *Anisakis*. However, elements involved in the molting, such as structural constituents of cuticle, different collagens and cuticlins, feature the list of differentially expressed genes in larvae infecting both rat and sea bass, with the distinction that those in rat have been increasingly upregulated, while in the sea bass, some were even strongly downregulated. As production of cuticle constituents is stage- and molt cycle-specific and precedes the molting by hours in *C. elegans* (Page and Johnston, 2007) or possibly even days in *A. suum* (Wang, 2014), we hypothesize that *A. pegreffii* larvae are in the initial stages of their molt process, more advanced in a rat than in a seabass. This is corroborated by the fact that some of key enzymes for collagen assembly did not show differential regulation in any host, such as prolyl-4 hydroxylase (P4H). However, protein disulfide isomerase 2 (PDI-2), essential for embryonic development, proper molting, extracellular matrix formation and normal function of P4H in nematodes (Winter et al, 2007; 2013), was found slightly upregulated in rat-infecting larvae.

According to other most represented KEGG pathways, biosynthesis of secondary metabolites and oxidative phosphorylation were positively perturbed in both hosts, although evidently stronger in rat, which reflects the need to satisfy various metabolic needs and increased demand for energy-rich molecules, such as ATP. Glycolysis/ gluconeogenesis-associated DEGs were noted in larvae from both hosts with fatty acid metabolism significant only for those infecting rat. This confirms that carbohydrate metabolism is constitutively the essential energy source for infecting larvae, but the fatty and amino acid metabolisms are required for infection of homeotherm host. Similarly, *A. suum* L3 found in lung also show a prominent upregulation of lipid/ fatty acid metabolism that eventually decreases with the transition to L4 and adults (Wang, 2014). Conversely, quiescent non-feeding L3 of the strongylid *Haemonchus contortus* that completes its lifecycle in the abomasum of ruminants, depend mainly on stored lipid reserves to survive adverse conditions in the pastures before reaching its next host (Laing et al, 2013). Fascinating studies from *C. elegans* confirm that monosaturated lipids regulate fat accumulation and longevity, saturated fatty acid acclimation to temperature, PUFAs are required for growth, reproduction, neurotransmission, and as precursors for signaling molecules (Watts and Ristow, 2017), indicating how complex these pathways are. Nonetheless, we suggest that the similarities between the profiles triggered in *A. suum* pig infection and *A. pegreffii* rat infection are likely correlated to homeothermy of the accidental host.

Upregulated elements of KEGG Microbial metabolism in diverse environments only in *Anisakis* rat-infecting larvae further supports the effort of the larvae to adapt to environmental and metabolic changes and survive stress conditions in the homeotherm host. This pattern consists of different metabolic processes, such as carbohydrate, carbon fixation, methane, nitrogen, sulphur, amino acid metabolism, as well as metabolism of cofactors and vitamins, and xenobiotic degradation effectuated by bacteria (https://www.genome.jp/kegg-bin/show_pathway?map01120), being usually expressed in response to the heat shock (Tripathy et al., 2014). Noteworthy is that the most upregulated element of this pathway in both hosts was mitochondrial glutamate dehydrogenase, a crucial enzyme linking nitrogen and carbon metabolism where ammonia is either assimilated to provide glutamate as nitrogen storage molecule, or dissimilated to alpha-ketoglutarate for the tricarboxylic acid (TCA) metabolism. Unfortunately, while its role to provide of reducing equivalents in form of NADPH required for downstream redox reactions essential in *Plasmodium falciparum* antioxidant machinery has been rebutted (Storm et al., 2011), scarce information exist for its function in helminths. More precisely, structural models give no tangible implication for its functional role in host infection, being also dismissed as unsuitable as a drug target (Brown et al., 2014). However, another highly expressed transcript during rat infection listed within this KEGG was enolase. It is a multifunctional glycolytic enzyme found engaged in adhesion and invasion of intracellular apicomplexan *Cryptosporidium parvum* (Mi et al., 2017), as well as activation of fybrinolytic agent plasmin in *Schistosoma mansoni* intravascular life stages, where apparently helps the trematode to maintain anti-coagulated environment (Figueiredo et al, 2015). Similarly, nematode *Trichinella spiralis* enolase binds the host’s plasminogen to activate the fibrinolytic system, degrades the extracellular matrix and promotes larval penetration of the tissue barrier during invasion (Jiang et al., 2019), which could be speculated for *Anisakis* enolase as well.

In spite of the evident cues for cell proliferation in rat-infecting L3 indicative of molting toward L4 stage, less represented KEGG pathways show downreagulation in both Cell senescence and Cell cycle. This apparently conflicting status; i.e. the blocking of transcripts programmed to arrests cell proliferation in response to different damaging stimuli (Muñoz-Espín Serrano 2014; Zumerle and Alimonti, 2020), contrasted by the blocking of transcripts that should support the cell cycle, could add to two likely coupled strategies. Firstly, KEGG Cell senescence and Cell cycle show similar levels of downregulation, implying that both processes are balanced, acting mutually in a feed-back loop. Being tightly coupled to cell growth, the efficient ribosynthesis enabled by high transcription rate of rDNA genes and the activity of ribosomal polymerases, allows a rapid cell proliferation required to meet cellular needs for ribosomes. However, under the stress conditions that affect the cell cycle and intracellular energy status (e.g. lack of nutrients), change in ribosynthesis is one of the cell strategies to retrieve homeostasis (Sengupta et al., 2010), likely to manifest in this case. Secondly, the homeothermic host environment, although offers the initial cues for growth, moulting and reproduction of *A. pegreffii* L3, possess additional conditions acting upon *Anisakis’* further development, which consequently result in parasites failure to survive. We can only speculate whether rat microbiome additionally contributes to such outcome, as in general microbiome interaction between the host and parasite differs for each specific case (Zaiss and Harris, 2016). However, it is logical to assume that *Anisakis* evolutionary has not been in contact with a terrestrial, homeotherm microbiome, which is likely to impose additional pressure on the larval survivor in the accidental host.

From the physiological stimuli-reaction standpoint, we can also hypothesise that the fate of the larval *Anisakis* in the accidental host is simply a result of larval exhaustion of nutrients and energy, spent during undetermined migration towards a niche that does not prove to be adequate, and stimulated by elevated host’s temperature. In contrast, in the parathenic exothermic host where metabolic pathways are moderately upregulated or silenced, larvae prepare for paratenesis that warrants their survival. This is inferred through FoxO signalling pathway, which was substantially downregulated in the accidental and upregulated in the paratenic host. While the pathway encompasses transcription factors that regulate expression of many downstream genes involved in cellular processes such as apoptosis, cell-cycle control, glucose metabolism, oxidative stress resistance, and longevity (Tia et al., 2018), the highest upregulated transcript (more then 10-fold) in sea bass-infecting *Anisakis* is gamma-aminobutyric acid receptor-associated protein (GABARAP), recognised as a hallmark for autophagy (Oshumi, 2014), but also encompassed within KEGG Longevity regulating pathway. GABARAP accumulates in the pericentriolar material under nutrient rich conditions, from which is translocated during starvation to form autophagosomes (Joachim et al., 2017). Although FoxO-GABARAP axis has been studied in colorectal and ovarian cancers, it is tempting to speculate that the autophagy in fish-infecting larvae represents a safety mechanism for their successful survival. Namely, in cancer FoxO3a senses variation in AMP/ATP ratio by decreased gycolysis, which activates FoxO3a transcriptional program, resulting in activation of genes involved in autophagic flux, namely GABARAP, GABARAPL1, GABARAPL2 and MAP1LC3 (Grossi et al., 2019). In general, autophagy is an essential cellular mechanism that enables the cell to counter-balance various demands by producing autophagosomes that engulf a wide range of intracellular material and transport it to lysosomes for subsequent degradation (Nakatogawa, 2020). While the basal autophagy acts as a housekeeping mechanism, the inducible autophagy starts by engulfment of bulk cytoplasm in times of stress, such as nutrient deprivation. Although this process still needs to be characterised in parasitic helminths, *C. elegans* employs it to remove aggregate-prone proteins, paternal mitochondria, spermatid-specific membranous organelles; remodeling during dauer development; degradation of the miRNA-induced silencing complex; synapse formation and in the germ line; to promote the stem cell proliferation; removal of apoptotic cell corpses; lipid homeostasis and in the ageing process (Palmisanoa and Meléndez, 2018). We suggest that in *Anisakis* larvae during exothermic conditions of infection, GABARAP induced through FoxO and/or Longevity regulating pathway, triggers autophagy (KEGG Autophagy-animal, -yeast, -other) that eventually balances the metabolic rate in larvae by clearing damaged/used organelles to prepare the nematode for indefinite paratenesis, necessary to counteract larval ageing and death. To further support this, KEGG Longevity regulating pathway showed to be upregulated in larvae infecting seabass and downreagulated in those infecting a rat. The relationship between *Anisakis* autophagy, metabolic balancing and longevity was supported by one of the highly expressed elements of KEGG Longevity regulating pathway - adiponectin receptor preotein 2; one of two transmembrane receptors that bind and activate adiponektin in humans, regulating glucose and lipid metabolism (Buechler et al., 2010). As the calorie restriction and consequent limited buildup of toxic cellular waste has been shown to extend the lifespan in a range of organisms (Mannack and Lane, 2015), it seems that the same strategy could be employed for *Anisakis* paratenesis.

### Anisakis pegreffii virulence factors and their potential drug-targeting

We defined *A. pegreffii* virulence factors as those transcripts expressing the highest upregulation common for *A. pegreffii* migrating through both hosts in respect to larvae that failed to do so. From the initial 65 transcripts, we selected 35 that showed at least two-fold change expressed in at least one of the host (Table 3), and then discarded constitutive cuticle elements and those with no annotation. That left us with several catalysts and transporters, some being recognised as excretory/ secretory (ES) products; cytosolic non-specific dipeptidase (CNDP2), leukotriene A-4 hydrolase (LKHA4), aspartic protease 6 (ASP6), ATP-binding cassette sub-family B member 9 (ABCB9), and UDP-glucuronosyltransferase (UGT3). Some of them (CNDP2, ASP6 and ABCB9) have subfamily/ subgroup members identified as surface-exposed molecules on the extracellular vesicles (EVs) of the trematode *Fasciola hepatica* (de la Torre-Escudero et al., 2018). EVs have been recognised as essential mediators of communication (through molecular signals such as proteins, lipids, complex carbohydrates, mRNA, microRNA and other non-coding RNA species) between parasite and host, particularly in helminth immunomodulatory strategy, suggesting that these transcripts might have a role in *Anisakis*-host cell signaling. Interestingly, none have been identified in previous works (Kim et al., 2018; Llorens et al., 2018), probably because those focused on transcriptomic profiles of dormant, non-infecting or *in vitro* cultured larvae. Specific temporal regulation of certain virulence factors might also depend on their other putative functions and co-expression networks. For instance, hyaluronidase, a hydrolytic enzyme that degrades the glycosaminoglycan hyaluronic acid, erected as crucial for pathogenesis of *Ancylostoma caninum*, *Anisakis simplex* and *A. suum* (Hotez et al, 1994; Wang, 2014; Ebner et al, 2018) showed no differential regulation in either of hosts studied herein, albeit expressed in the transcriptome. Rhoads et al. (2001) noted its release in *A. suum* when transitioning from L3 to L4, corroborating its other important functions next to facilitation of larval penetration, such as feeding, proper molting and development. If this is also true for *Anisakis*, we might have missed the point when the need for this enzyme surges.

The versatile gene content and unequal protein family representation among nematodes and other helminths has been established and reflects uniqueness of parasite biology and different pathogenic strategies (Coghlan et al, 2019). It is also the result of specificities of each host-parasite relationship formed during their evolution. In order to investigate whether virulence factors erected for *A. pegreffii* are shared between other helminths and investigate their potential for repurposing of existing therapeutics, we have performed orthology-directed phylogenetic analyses of *A. pegreffii* ABCB9, UGT3, ASP6, LKHA4 and CNDP2 to visualize gene gain and loss among representative helminths (Figure 4). We have also determined and identified the 3-dimensional structures and catalytic sites for all five above mentioned DEGs utilizing online modelling techniques and comparing our models to structures co-crystalized with inhibitors or substrate analogues (Figure 5).

The results of our phylogenetic analyses support the premise that the four families (ABCB9s, ASP6s, LKHA4s and CNDP2s) are present in almost all Nematoda, platyhelminthes and metazoans examined, except for the Trematoda, Cestoda and vertebrate species that have lost UGT3. The high levels of duplication and wide-spread occurrence of all five target genes in closely related *T. canis, A. suum, P. univalens* and also *H. contortus*, suggests that these genes may have vital biological functions as virulence factors in these extant species. However, it is not clear why the Trematoda, Cestoda and outgroup species examined do not contain any recognizable UGT3 gene, but leads us to propose that these observations may be due to substantial divergence of UGT3 or incorporation of protein domains by horizontal gene transfer that has not been detected in this study.

Aspartic protease 6 is an endopeptidase involved in haemoglobin digestion, tissue penetration or host-derived nutrient digestion in helminths (Koehler et al., 2007; Ebner et al. 2018). The importance of proteases and protease inhibitors for helminths is reflected in their vast representation in nematode and platyhelminth species, as noted in a large comparative genomic study of parasitic worms (Coghlan et al, 2019). Aspartic proteases have been found especially abundant in Clade IV and V Nematoda, which is in general agreement with our phylogenetic inference of orthologues. *Asp6* is one of the first transcripts upregulated in *Ascaris suum* upon contact with porcine epithelial cells (Ebner et al, 2018) and an altered homologue of aspartic protease 1 from *Necator americanus* has been selected as a target for human hookworm vaccine development (Hotez et al, 2013). Aspartic protease 6 has also been targeted in trypanosamotids therapy (causative agents of leishmaniasis, Chagas’ disease and sleeping sickness) by canonical (DAN, EPNP, pepstatin A) and anti-HIV aspartic peptidase inhibitors (amprenavir, indinavir, lopinavir, nelfinavir, ritonavir and saquinavir). The latter inhibitors affected parasite’s homeostasis, through elevated production of reactive oxygen species, apoptosis, loss of the motility and arrest of proliferation/growth (Santos et al., 2013). Drugbank lists several experimental chemotherapeutics targeting aspartic proteases of malarian parasite *Plasmodium falciparum* (artenimol) and fungus *Candida albicans* (ethylaminobenzylmethylcarbonyl, 1-amino-1-carbonyl pentane, butylamine, 1-hydroxy-2-amino-3-cyclohexylpropane, 4-methylpiperazin-1-Yl carbonyl), as well as other human targets with aspartic-type endopeptidase activity involved in other pathogenesis, such as in Alzheimer’s disease (https://go.drugbank.com).

ATP-binding cassette sub-family B member 9 belongs to a large group of multidrug resistance (MDR)/ transporters associated with antigen processing (TAP) transmembrane proteins. ABCB9 proteins are known therapeutic targets in disease (Shintre 2013) and well known to bind and confer drug resistance in cancer cells (Jin 2012). Located in lysosomal membrane, they use ATP-generated energy to translocate cytosolic peptides to the lysosome for processing (Zhao et al., 2008). However, until their functional characterisation in helminths, we cannot state whether the protein is lysosomal, or rather involved in the efflux of chemically unchanged organic xenobiotics, as some of ABCB members. The efficiency of *in vitro* anisakiasis treatment by inhibitors of other MDR members (ABCB/P-glycoprotein and MRX1) proved to be efficient (Mladineo et al., 2017), but no treatments targeting ABCB9 so far have been reported (https://go.drugbank.com). A genome-wide identification of ABC transporters in monogeneans identified orthologues of ABCB family in *Gyrodactylus salaris, Protopolystoma xenopodis, Neobenedenia melleni*,and specifically ABCB9 in *Eudiplozoon nipponicum*, as well as *C. elegans* (Caña-Bozada et al., 2019). This is in general agreement with our orthology inference, except for the presence of orthologues of ABCB9 in *G. salaris*, which might be the result of stringent criteria we used for orthologue identification. In general, ABC transporters show independent losses and expansions within parasitic worms (Coghlan et al, 2019).

Cytosolic non-specific dipeptidase, also known as carboxypeptidase of glutamate-like (CPGL), catalyses the hydrolysis of peptides, being found significantly upregulated in adult stages of a tapeworm *Taenia pisiformis*, presumably associated to amino-acid transport and metabolism (Zhang, 2019). However, human CNDP2 figures as an important tumor suppressors in gastric, hepatocellular and pancreatic cancers that inhibits cell proliferation, and induces apoptosis and cell cycle arrest, via activation of mitogen-activated protein kinase (MAPK) pathway (Zhang et al., 2013). It is tempting to speculate whether *Anisakis* CNDP2 also serves in MAPK pathway, the latter employed in cell communication during helminth development and homeostasis (Dissous et al., 2006). No treatments targeting CNDP2 so far have been reported in Drugbank (https://go.drugbank.com).

Leukotriene A4 hydrolase has been studied as ESP of *Schistosoma japonicum*. The trematode synthesise proinflammatory mediators prostaglandins through arachidonic-acid metabolism that uses lecithin to generate arachidonate, converts it in leukotriene A4 and then more stable leukotriene B4 by leukotriene A4 hydrolase (The Schistosoma japonicum Genome Sequencing and Functional Analyses Consortium, 2009). Interestingly, prostaglandins and leukotrienes contribute to metabolism or maturation of the organism and communication with the host on a cellular basis, acting as immunomodulators and eosinophil attractants (Noverr et al, 2003). In *S. japonicum* they have been suggested to induce chemokine-receptor-mediated cell migration and leukocyte migration into inflamed tissue, which for the parasite is essential for survival as it promotes granuloma formation around expelled eggs. How *Anisakis* larvae benefit from stimulation of proinflammatory host reaction through upregulation of leukotriene A4 hydrolase is not clear, as inflammation favors propagation of only few parasites (Sorci and Faivre, 2009). Of 22 investigated drugs targeting human LKHA4, only captopril has been approved as an inhibitor of angiotensin-converting enzyme in regulation of blood pressure, having also an unknown pharamcological action on LKHA4 (https://go.drugbank.com).

*Anisakis* UDP-glucuronosyltransferase (UGT) homologues were significantly upregulated in larvae penetrating epithelial barriers of both hosts types in our study (UGT47, UGT50, UGT58), and a couple more reconstructed in the transcriptome were classified as DEGs only in rat (upregulated UGT50 and downregulated UGT60). UDP-glucuronosyltransferases catalyze glucuronidation reaction, the addition of polar glucuronic acid to lipophilic substrates promoting their elimination and clearance from the organism. As such they are important part of detoxification process and have been associated with benzimidazole resistance phenotype in *H. contortus* (Matoušková et al, 2018) or naphtalophos biotransformation (Kotze et al., 2014). Thirty four UGTs were reconstructed in the transcriptome of *H. contortus* (Laing et al, 2013), with four of these are putative orthologues to *A. pegreffii* UGT3 (Figure 4). According to KEGG orthology, UGTs interlink various metabolic pathways, such as Pentose and glucuronate interconversions, Ascorbate and aldarate metabolism, Steroid hormone biosynthesis, Retinol metabolism, Porphyrin and chlorophyll metabolism, Metabolism of xenobiotics by cytochrome P, Drug metabolism, Biosynthesis of secondary metabolites. Because it is essential for adult worm survival and due to its large, easily-targeted extracellular domain, in *Brugia malayi* it was selected as a potential drug target for lymphatic filariasis (Flynn et al, 2019). Two compounds targeting UGT3 are experimental and/or under investigation; kaempherol and quercetin, both flavonols with unknown pharmacological action and antioxidant properties. The latter however, has many targets in addition to UGT3, but it mainly inhibits quinone reductase 2 of the *P. falciparum* causing the lethal oxidative stress (https://go.drugbank.com). Interestingly, UGTs are also known for mediating metabolic inactivation of lipophilic cancer drugs. However, recently has been suggested that dysregulation of UGT expression might promote oncogenic pathways by metabolizing endogenous molecules such as steroids and bioactive lipids and disturbing homeostasis (Allain et al, 2020). The versatile role of these enzymes and detoxification processes is also observed in dauer *C. elegans* larvae where it has been proposed that cytochrome P450, UDP-glucuronosyltransferase and glutathione S-transferase perform vital clearance of toxic lipophilic and electrophilic metabolites associated with ageing and reduced longevity (McElwee et al, 2004). In study performed by Rausch et al (2018), UGTs and glutathione S-transferases were differentially regulated between *Heligmosomoides polygyrus* infecting germfree and conventional specific pathogen-free mice, suggesting that these detoxification enzymes also participate in nematode sensing of its microbial environment. Host microbiome has been demonstrated as important factor in shaping host-parasite relationships (Zaiss and Harris, 2016); furthermore *H. polygyrus* showed reduced fitness in germfree mice (Rausch et al, 2018). This represents an axis of investigation that must be explored in the future for *A. pegreffii* and might prove to be the missing explanatory link behind its success or failure to infect evolutionary distant hosts.

The discovery of new anthelmintic drug targets with broad-spectrum efficacy is expensive and time consuming. At present, approximately $2.6USD billion in direct costs and 10-15 years is the average length of time required to progress from the concept of a new therapy or drug target to a new molecular entity on the market Tamimi and Ellis. 2009). In addition, the rate of success is less than 5%, with less than 20% of compounds entering clinical trials actually receiving FDA approval over this time (Kola and Landis 2004; DiMasi et al., 2010). Our tertiary structure predictions and modelling analyses present the bases for the repurposing of selected candidate and currently available inhibitor molecules that should be incorporated in future investigations and may provide broad-spectrum efficacy particularly for all Clade III and V nematodes examined. For instance, BES is a well-known dipeptidase inhibitor for the aforementioned enzyme classes 3.3.2.6 and 3.4.13.18 (Andberg et al., 2000; Lenney 1990), while the ATP nucleotide analogue known as ACP would most likely bind ABCB9 due to the highly homologous nature of the active sites. Overall, inhibition may be possible with these candidate molecules, but more importantly targeted drug discovery efforts that could produce highly selective nematode species-specific inhibitors, might benefit from utilizing the drug sites modelled herein.

## Table Legends

**Table 1.** Summary of the design and outcome of experimental infections of *Dicentrarchus labrax* and *Rattus norvegicus* with *Anisakis pegreffii* L3 larvae. Number of intubated animals, sampling time, minimum, maximum and median percent (%) recovery rate, larval phenotype and site of collection within the host are shown.

**Table 2.** Statistics of *Anisakis pegreffii* de novo assembly and annotation summary.

**Table 3.** Differentially expressed genes (DEGs) identified in *Anisakis pegreffii* L3 migrating vs non-migrating larvae during experimental infections of a paratenic and an accidental host, *Dicentrarchus labrax* and *Rattus norvegicus*, respectively. Top 10 DEGs show different regulation in the two hosts and bottom 35 show congruent profiles for both hosts and at least two-fold change in expression in one of the hosts. Putative virulence factors selected as potential drug targets are depicted in bold. DEGs were identified at Benjamini-Hochberg false discovery rate (FDR) < 0.05.

## Supplementary material

**Supplementary Figure 1.** Frequency distribution of differentially expressed genes (DEGs) in top 10 Gene Ontology terms in Biological process, Molecular function and Cellular component identified in *Anisakis pegreffii* L3 larvae during experimental infections of a paratenic and an accidental host, *Dicentrarchus labrax* and *Rattus norvegicus*, respectively.

**Supplementary Table 1.** Sample description and formation of RNAseq libraries.

**Supplementary Table 2.** The reference assembled transcriptome of *Ansiakis pegreffii* with nucleotide and amino acid sequences and functional annotation.

**Supplementary Table 3.** Gene-level raw counts of each RNAseq library used for differential expression analyses of *Anisakis pegreffii* L3 larvae during experimental infections of *Dicentrarchus labrax* and *Rattus norvegicus*.

**Supplementary Table 4.** Complete list of differentially expressed genes (DEGs) identified in *Anisakis pegreffii* L3 larvae during experimental infections: Migrating vs. non-migrating larvae in rat *Rattus norvegicus*, migrating vs. non-migrating larvae in seabass *Dicentrarchus labrax* and the interaction: larvae showing different regulation in two hosts.

**Supplementary Table 5.** Results of enrichment analyses of GO terms within sets of DEGs identified in *Anisakis pegreffii* L3 larvae during experimental infections: Migrating vs. non-migrating larvae in rat *Rattus norvegicus*, migrating vs. non-migrating larvae in seabass *Dicentrarchus labrax* and the interaction: larvae showing different regulation in two hosts. Goseq package for R was used for the analyses taking gene length bias into account.

**Supplementary Table 6.** Results of enrichment analyses of KEGG metabolic and signaling pathways via KEGG Orthologues (KO) within sets of DEGs identified in *Anisakis pegreffii* L3 larvae during experimental infections: Migrating vs. non-migrating larvae in rat *Rattus norvegicus*, migrating vs. non-migrating larvae in seabass *Dicentrarchus labrax* and the interaction: larvae showing different regulation in two hosts. Goseq package for R was used for the analyses taking gene length bias into account.

**Supplementary Table 7.** Complete list of orthologous protein families for each of the five DEGs of interest across 28 species including additional annotations.

## Acknowledgements

High performance computing was conducted using the resources of computational cluster Isabella hosted by the University Computing Centre, University of Zagreb (SRCE), Croatia.

